# epiAneufinder: identifying copy number variations from single-cell ATAC-seq data

**DOI:** 10.1101/2022.04.03.485795

**Authors:** Akshaya Ramakrishnan, Aikaterini Symeonidi, Patrick Hanel, Michael Schubert, Maria Colomé-Tatché

## Abstract

Single-cell open chromatin profiling via the single-cell Assay for Transposase-Accessible Chromatin using sequencing (scATAC-seq) assay has become a mainstream measurement of open chromatin in single-cells. Here we present a novel algorithm, epiAneufinder, that exploits the read count information from scATAC-seq data to extract genome-wide copy number variations (CNVs) for individual cells, allowing to explore the CNV heterogeneity present in a sample at the single-cell level. Using different cancer scATAC-seq datasets, we show how epiAneufinder can identify intratumor clonal heterogeneity in populations of single cells based on their CNV profiles. These profiles are concordant with the ones inferred from single-cell whole genome sequencing data for the same samples. epiAneufinder allows the addition of single-cell CNV information to scATAC-seq data, without the need of additional experiments, unlocking a layer of genomic variation which is otherwise unexplored.

## Introduction

Aneuploidy and copy number variations describe DNA duplication and deletion events that range from a small number of base pairs in the genome to whole chromosomes. Both conditions have been involved in disease, and are especially prominent in cancer^1^. In fact, more than ∼90% of solid tumors are aneuploid^2^, leading to the hypothesis that aneuploidy confers a growth advantage to cancer cells^3^. However, the relationship between aneuploidy and tumorigenesis remains not clearly understood^4,5^. Meanwhile, aneuploidy is highly detrimental to normal cell development and growth^6,7^.

Because of its biological and clinical relevance, aneuploidy is widely studied using different experimental strategies^8^. Traditionally, CNVs have been studied with spectral karyotyping or (interphase) fluorescence in situ hybridization (FISH), which are single-cell low throughput methods that lack precise genomic resolution^8^. On the contrary, comparative genomic hybridization (CGH), whole exome sequencing (WES) or whole genome sequencing (WGS) provide high genomic resolution but do so at the expense of losing single-cell resolution^8^. Single-cell whole genome sequencing (scWGS) offers a compromise, interrogating copy number gains and losses in a high throughput fashion at the single-cell level and at a higher genomic resolution than FISH or spectral karyotyping^8–10^. Despite being considered the gold-standard ground-truth for the quantification of CNV heterogeneity in large single-cell populations^9^, scWGS is not often used in the laboratory compared to other single-cell sequencing techniques. Attempts have been made to call copy number variations from single-cell gene expression data, however, at high genomic resolution these are confounded by physiological variation in expression levels^11^, and calling CNVs from single-cell gene expression alone proves challenging^12–16^.

A novel single-cell measurement technique, single-cell Assay for Transposase-Accessible Chromatin using sequencing (scATAC-seq), has become a new mainstream measurement in single-cells^17^, partially owing to its implementation in the 10x platform^18^. Single-cell chromatin openness measurements require the sequencing of the DNA of the cell, instead of the RNA like in scRNA-seq, hence scATAC-seq measurements have the potential to better recapitulate the DNA content from single-cells. However, scATAC-seq data is extremely sparse, making it challenging to directly extract copy number calls from the data^18,19^.

In this paper we present a new algorithm, epiAneufinder, that calls copy number variations at the single-cell level from scATAC-seq data. epiAneufinder uses binary segmentation combined with an appropriate choice of distance measure to identify putative breakpoints in the genome, and subsequently calls gains and losses per identified segment. epiAneufinder can identify single-cell CNVs from scATAC-seq data alone, without the need of a reference euploid sample and without the need of supplementing the data with other data modalities. We demonstrate the performance of epiAneufinder by applying it to different scATAC-seq cancer datasets for which an orthogonal measurement of CNVs is available (either scWGS or WES). In conclusion, epiAneufinder allows the addition of an extra level of genetic information, namely CNVs, to scATAC-seq or single-cell multiome (combined scRNA and scATAC-seq) data, without the need of any extra experimental effort. epiAneufinder is available as an R package at https://github.com/colomemaria/epiAneufinder.

## Results

### epiAneufinder algorithm

#### Genome binning

The goal of epiAneufinder is to segment the genome into regions of gain, loss, and normal copy number per single-cell. To do that, epiAneufinder uses the number of reads from scATAC-seq data mapping to a genomic region as a proxy of the number of DNA copies present in that region, for every single-cell. To overcome the coverage sparsity inherent to single-cell sequencing, lowly covered cells are filtered out, the genome is binned into equally sized windows (by default, window size is 100,000 bp) and the number of mapped reads per window are quantified (Fig. 1a). Furthermore, we remove the ENCODE blacklisted set of regions^20^, comprised of certain genomic locations, such as telomeric ends and repetitive regions, that have systematic biases in their mappability and that would therefore bias the copy number inference. For every dataset, epiAneufinder also removes bins that have zero counts in >85% of all the cells, to discard genomic areas that have low mappability in every dataset specifically. All parameters can be adjusted by the user.

**Figure1:**
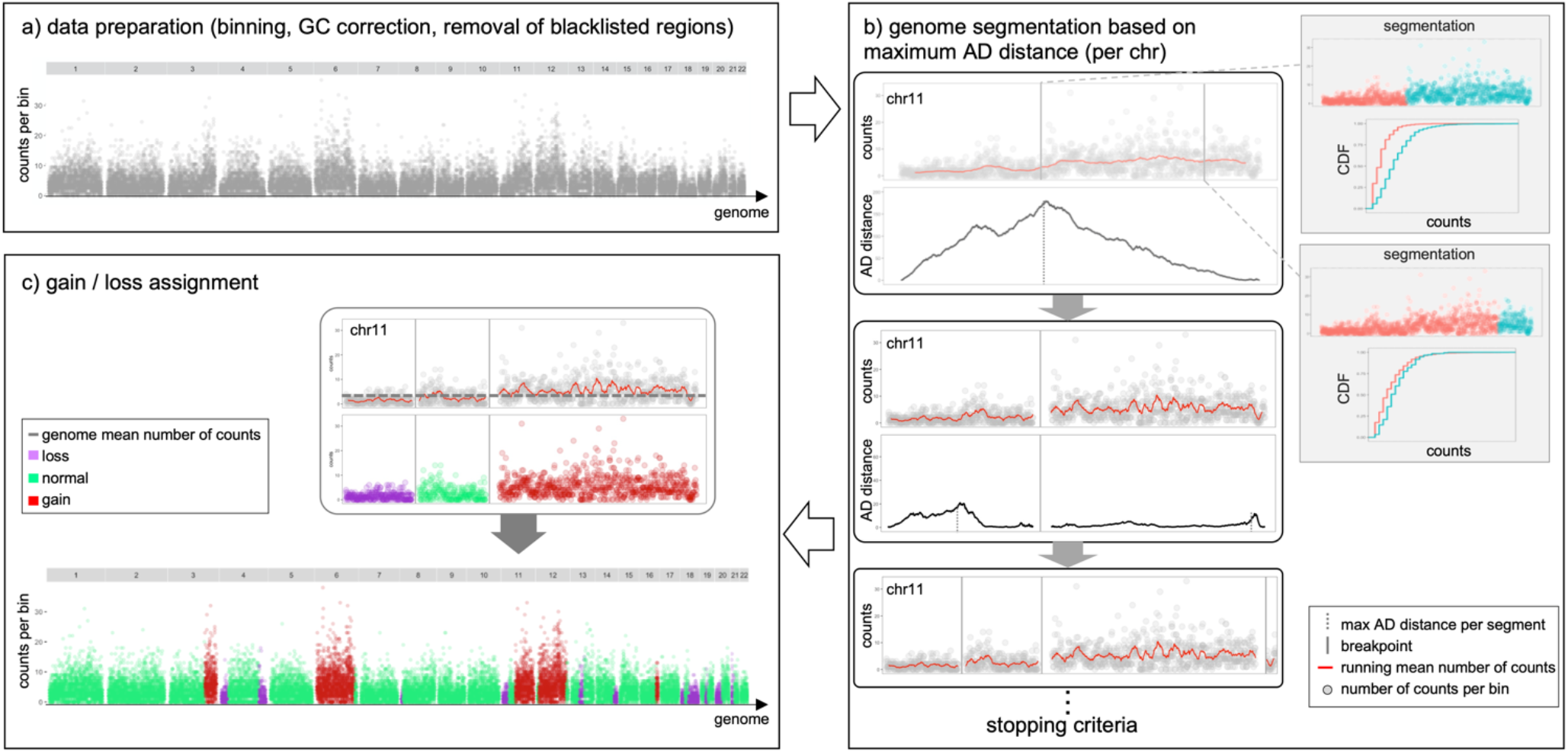
epiAneufinder algorithm: a) The genome is binned (default 100,000 bp windows) and the number of reads in every bin are quantified. Blacklisted regions are removed and the data is GC corrected. Bins with zero counts in >85% of all the cells are removed. b) Binary segmentation is applied iteratively per chromosome by computing the AD distance between the left (red) and right (turquoise) count distribution (CDF=cumulative density function). In every segment the position with the highest AD distance is considered a breakpoint until a stopping criteria is reached. c) After breakpoints have been identified per chromosome, every segment is assigned to the state loss, normal or gain based on the read-count fold change over the genome-wide mean.

After that, the binned dataset is GC corrected using a LOESS fit. The correction factor is obtained by fitting the raw read counts to the GC content per bin:

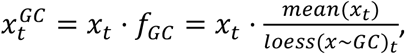

where *x*_*t*_ is the number of reads in bin *t, mean*(*x*_*t*_) is the average read count per bin and *GC* is the percentage of base pairs which are GC per bin.

#### Binary segmentation and copy number calling

After data preparation, epiAneufinder applies a binary segmentation algorithm to each single-cell separately. Binary segmentation is a technique used to detect change points in signals by identifying positions of data distribution changes. In the case of CNV calling, the assumption is that the distribution of reads mapping per bin is different for a gained, a lost, and a normal copy number region (SI Fig. 1). The goal is to identify the positions in the genome where the distribution changes take place. To do that, the algorithm scans the genome of every single-cell per chromosome, and calculates the Anderson-Darling (AD) distance (*d*_*AD*_) between the read distributions at the left and right of every bin (Fig. 1b). The AD test is a non-parametric test that measures agreement between distributions. It was initially developed to check for normality^21^, but it was subsequently modified^22^ to measure the distance between any two empirical distributions:

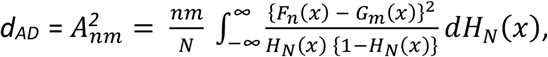

where *F*_*n*_(*x*) and *G*_*m*_(*x*) are the read distribution functions for the two genomic segments to compare (with lengths *n* and *m*, respectively), and *H*_*N*_(*x*) is the distribution function of the combined segments: *H*_*n*_(*x*) = {*nF*_*n*_(*x*) + *m*G_*m*_(*x*)}/*N* with *N* = *n* + *m*. It was chosen here to detect copy number differences because it emphasizes the difference between distribution tails.

The position in every chromosome that maximizes the AD distance is kept as the most likely breakpoint (Fig. 1b). The same procedure is then repeated iteratively on the two resulting segments, until a total number of breakpoints are identified per chromosome (by default seven) (Fig. 1b). After all the breakpoints have been identified genome-wide, epiAneufinder prunes out breakpoints with an AD distance lower than the genome-wide mean.

Finally, every segment is assigned to the state gain, loss, or normal copy number. To do that, epiAneufinder calculates the trimmed-average number of reads per identified segment, defined as the average number of reads per bin in every segment without the lowest 0.1 and highest 0.1 quantile bins (these values can be adjusted by the user). The segments with a trimmed-average number of reads with z-score between [-1, 1] are assigned to the genome-wide mean. Then, for every segment, the algorithm calculates its rounded integer fold change over the genome-wide average number of reads. For a value of 0 the segment is assigned to the state “loss”, for a value of 1 to the state “normal” and for a value >=2 to the state “gain” (Fig. 1c). For visual representation and to identify CNV clones, the single-cells are clustered based on their copy number profiles using Euclidean distance and Ward Clustering.

In summary, epiAneufinder takes scATAC-seq BAM files or 10x fragment files as input, and outputs a RDS file and a TSV file with the identified copy number states for each bin per cell, labeled as “loss” (0), “normal” (1) or “gain” (2). Other intermediate result files are also provided, such as the (GC corrected) binned number of reads per cell (RDS file), and the identified breakpoints per cell and per chromosome with their associated AD distance (RDS file). Moreover, plotting functions are available via epiAneufinder, to plot the resulting single-cell karyotypes and the clustering results. Run times for all the datasets presented are available in Table S1.

### Applications

#### epiAneufinder CNVs are concordant with the ones obtained from scWGS in a gastric cancer cell line

To demonstrate the performance of epiAneufinder, we analysed a recent scATAC-seq dataset from Wu et al.^19^. This dataset contains ∼3,500 aneuploid single-cells from the gastric adenocarcinoma cell line SNU601 with a mean coverage of 75,013 fragments per cell. The SNU601 cell line is known to contain a complex subclonal structure with multiple clones that harbor different copy number calls^23^. The same cell line was profiled using scWGS^23^, a measurement which can be used as an independent ground-truth for comparing the identified scATAC-seq CNV calls to.

We segmented the SNU601 genome into 100,000 bp windows and we quantified the number of reads per cell in every window. epiAneufinder was applied to every cell in the population with standard parameters, breakpoints were identified, and every segment was assigned to the state “gain”, “loss” or “normal”. We identified several clones in the dataset (Fig. 2a, SI Fig. 2). The main source of variation was the presence (clone 1) or absence (all other clones) of a whole chromosome gain in chromosome 6. Clone 1 was further split into two subclones (clones 1.1 and 1.2), mainly differentiated by the presence or absence of a gain in chromosome 12. Clone 2 was found to be nearly disomic. The two other clones could be split into different minor groups, the major source of variation being the presence of a longer or shorter gain in chromosome 3, as well as the presence of gains in chromosomes 11 or 12 and losses in chromosome 18.

**Figure2:**
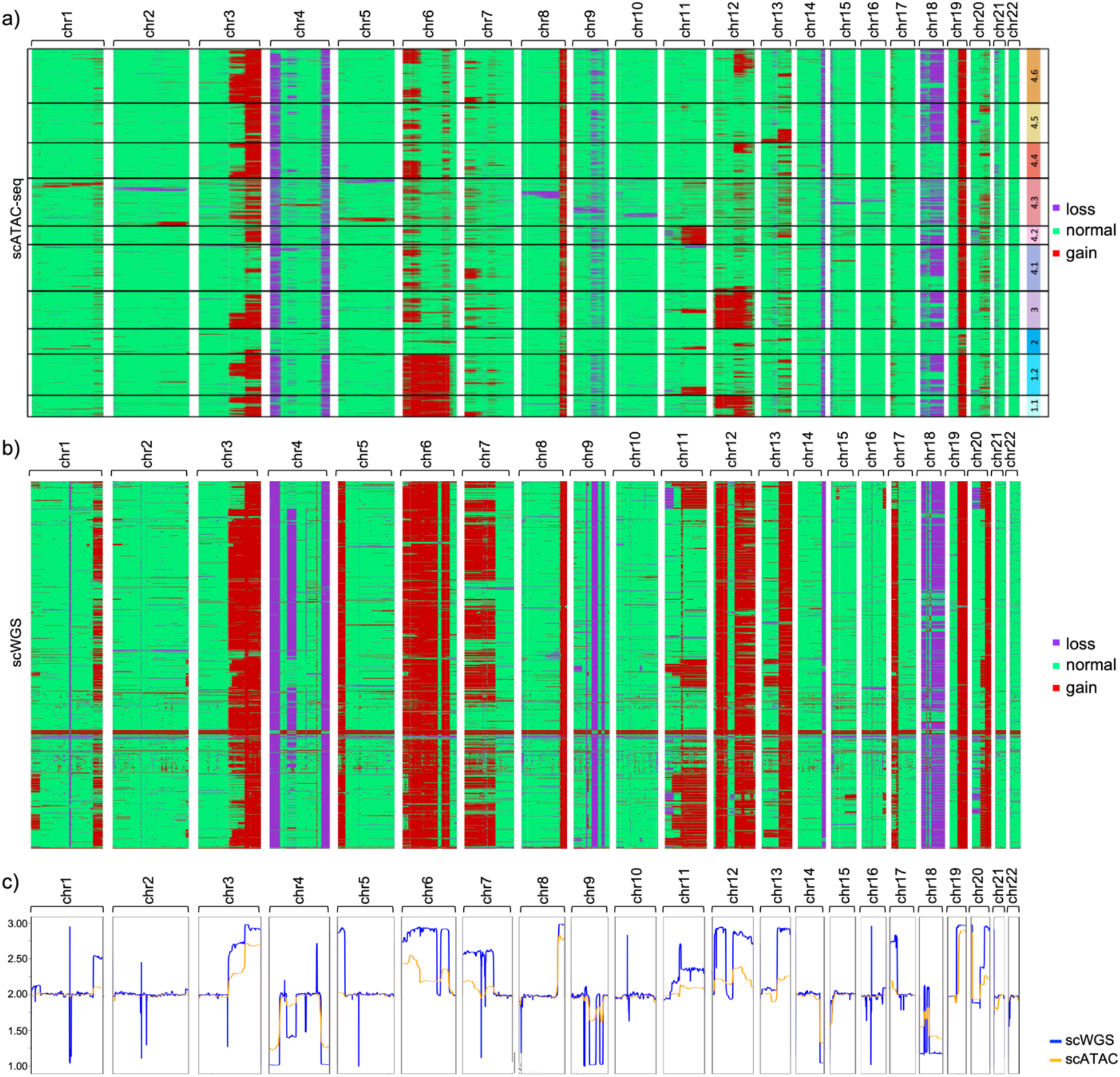
SNU601 cell line copy number variations: a) CNVs for the SNU601 cell line obtained from scATAC-seq data. Every row is a cell and every column is a chromosome. Karyotype clones are indicated in black boxes and numbered. b) CNVs for the SNU601 cell line obtained from scWGS data. Every row is a cell and every column is a chromosome. c) Pseudo-bulk CNV profiles for the DNA and ATAC data.

To validate our results, we analyzed a scWGS dataset for the same cell line, with 1,531 cells (average coverage of 707,188 reads per cell)^23^. Copy number gains and losses were called using the package aneufinder, designed to work with single-cell DNA sequencing data^24^, using the same 100,000bp window size as for the scATAC-seq. Despite the fact that the two datasets were produced by different laboratories using different techniques, the scWGS data presented a very similar CNV profile as the scATAC-seq dataset (Fig. 2b, SI Fig. 3).

To quantify the similarities between the two copy number profiles, we constructed the in-silico pseudo bulk copy number profiles, calculated as the mean of the gains and losses in every bin in the population. In general, the same pseudo-bulk gain and loss profile was observed for both modalities (Fig. 2c), and the pseudo-bulk DNA-seq and ATAC-seq profiles were highly correlated (Pearson correlation value 0.86 genome-wide (p<1e-307)). Considering the DNA-seq profile as the observed (true) value, and the scATAC-seq profile as the predicted one, the mean square error (MSE) of the detected scATAC-seq CNVs was computed, which was as low as 0.088 genome-wide (Table S2).

Generally, the pseudo-bulk scATAC-seq copy number profile tended to be less penetrant than the scWGS one, an effect that was observed both for gains and for losses (Fig. 2c). The scWGS dataset presented some sharp singularities around the centromeres/pericentromeres that were not detected on the scATAC-seq dataset; while a very penetrant gain on chromosome 5 observed on the scWGS dataset was not recovered in the scATAC-seq dataset (Fig. 2c). These discrepancies could be due to the stark differences in experimental protocols between scWGS and scATAC-seq measurements, combined with the fact that the DNA and ATAC experiments were performed independently by different laboratories(Wu et al 2021^19^ and Andor et al. 2020^23^), and it has been documented that despite its assumed homogeneity, cell lines can be highly variable^25^.

#### epiAneufinder uncovers CNV tumor heterogeneity in primary patient samples

We further applied epiAneufinder to two patient samples of basal cell carcinoma that were profiled using scATAC-seq^18^ and for which whole exome sequencing (WES) data had previously been generated^26^. For the scATAC-seq dataset, 2,040 and 504 cells were profiled for two patients (named SU006 and SU008), respectively, with an average number of fragments of 58,055 and 63,060. The genome of both samples was binned into 100,000 bp bins and epiAneufinder was applied with standard parameters to identify breakpoints and assign copy number gains and losses per cell.

Several clonal gains and losses were identified in the genomes of both patients (Fig. 3a, SI Fig. 4-5). Patient SU006 presented three main clones. One of them, clone 1, showed a low number of copy number aberrations. Clone 2 had two main subclones, one characterized by a whole chromosome 4 loss, and the other with losses in chromosomes 2 and 9, and a gain in chromosome 13. Clone 3 had several subclones characterized by gains of different lengths in chromosome 13, gains in chromosome 6 and losses in chromosome 9. Patient SU008 also presented a nearly-disomic clone (clone 2), as well as a clone mainly characterized by different combinations of gains in chromosomes 1, 3 and 6 (clone 1), and a clone with combinations of a loss in chromosome 13 and gains in chromosomes 2, 6 and 19 (clone 3). We compared epiAneufinder’s copy number profiles to the WES measurements for the same tumors^26^, which represented an orthogonal measurement of the average copy number gains and losses in the population. For both patients, the pseudo-bulk scATAC-seq CNVs were correlated to the WES profiles (Pearson correlation mean value of 0.53 and 0.56 for SU006 and SU008 (p<0.00001)) (SI Fig. 6). This level of agreement is notable given the sharp differences between the two data types and the different areas of the genome that they cover.

**Figure3:**
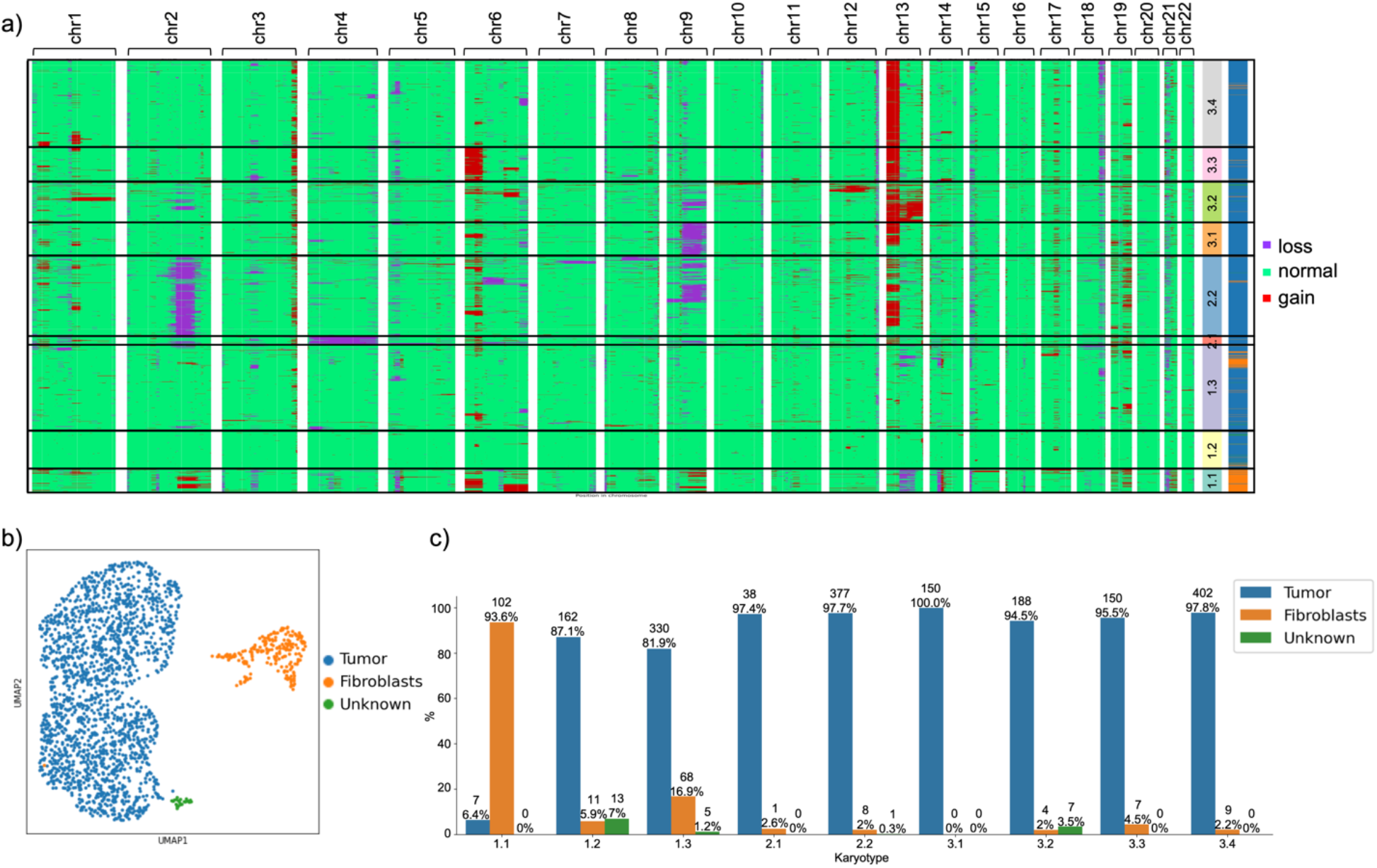
Copy number variations in a primary patient sample: a) CNVs for the SU006 primary sample obtained from scATAC-seq data. Every row is a cell and every column is a chromosome. Karyotype clones are indicated in black boxes and numbered. Cell types are indicated per cell in the side bar. b) UMAP embedding of the same cells showing tumor and fibroblasts. c) Correspondence between karyotype clones from (a) and cell types from (b).

Embedding the same cells based on their genome-wide scATAC-seq peak profile followed by Leiden clustering identified several clusters in both datasets, corresponding to tumor cells, fibroblasts and endothelial cells (Fig. 3b and SI Fig. 7). We compared the Leiden clusters and the karyotype clones (Fig. 3c and SI Fig. 8). For patients SU006 and SU008, fibroblasts corresponded mainly to the karyotype clones 1.1-1.3 and 3.1-3.2-3.3.2 respectively, which shared similar CNV profiles (Fig. 3 and SI Fig. 8). Endothelial cells were split into two karyotype clones in patient SU008: clone 2 (nearly diploid) and the subclone 3.3.3 (SI Fig. 8). The tumor cells showed very different characteristics depending on the patient of origin: while for patient SU006 the tumor was highly heterogeneous, containing 8 distinct karyotype clones (Fig. 3c and SI Fig. 8), in patient SU008 the tumor cells corresponded mainly to two similar clones (SI Fig. 8). The different CNV clones present in each tumor could not have been identified based on the embedding results (SI Fig. 8), emphasizing the relevance of epiAneufinder for discovering new sources of variation in the data. In both patients, the mostly-diploid karyotype clones (1.2 and 1.3 for SU006 and 2 for SU008) contained a mixture of all cell types (Fig. 3c and SI Fig. 8).

These results highlight how the single-cell CNV clones identified by epiAneufinder correspond to the cell types in the population, and exemplify the power of epiAneufinder to identify previously unknown distinct clones of tumor cells by their copy number profiles.

#### CNVs identification is marginally affected by sequencing read depth

To assess the robustness of epiAneufinder’s gain and loss calls to sequencing depth, we performed a simulation study. The SNU601 cell line dataset^19^ was downsampled from 100% coverage to only 20% of the initial reads (Tables S3-4), and used epiAneufinder with standard parameters to identify gains and losses in the population. The results of this simulation showed that, even when only 20% of the original number of reads were retained, on average over all the cells, 93% of the total genome remained in the same copy number state as identified in the fully covered dataset (Fig. 4a). Since the majority of the genome was in the normal state, we also investigated the robustness of copy number losses and gains calls separately upon downsampling. Percentage-wise, the identification of copy number losses was less affected by the downsampling than the identification of copy number gains: when considering only 50% of the initial coverage, ∼19% of bins with a copy number loss in the fully covered dataset were not identified as a loss any more (lost losses), compared to ∼23% of gains that were not identified any longer (lost gains) (Fig. 4c-d). As expected, the normal calls were the least affected (only 4.5% of the normal bins changed state at 20% downsampling) (Fig. 4b).

**Figure4:**
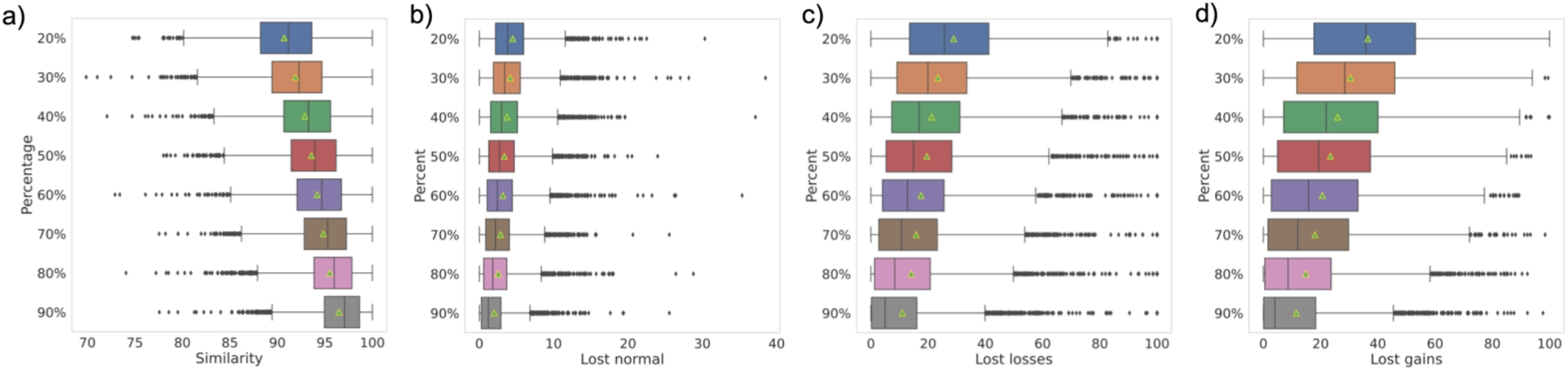
Simulation downsampling: a) percent of genome which is unchanged. b) Percent of normal calls, c) loss calls and d) gain calls which are lost upon downsampling.

Considering the fully covered dataset as the ground truth, we computed precision, recall and F1 scores for every state (Table S5). Precision, recall and F1 score was always >0.93 for all downsampling datasets for the normal state, indicating that overall the normal state was very robustly recovered regardless of sequencing depth. For the “gain” state, the lowest precision and recall were ∼0.8 and ∼0.6 respectively (at 20% coverage), meaning that there were fewer false positives than false negatives. False positives here are defined as new “gain” calls in the downsampled dataset in bins that were in a state “normal” or “loss” in the full dataset. Meanwhile, false negatives are bins that have lost their “gain” state in the downsampled dataset, to become either “normal” or “loss”. The trend was reversed for the “loss” state (precision and recall were ∼0.64 and ∼0.71 respectively, at 20% coverage). The number of false positives was higher than false negatives here, meaning that there were more novel “loss” calls in the downsampled dataset compared to the fully covered dataset.

Overall these results indicate that the genome-wide copy number gains and losses remained fairly stable upon downsampling even at low sequencing depths. The observed changes were mainly driven by “gain” states which were no longer identified in the low coverage datasets, as well as false positive “loss” calls stemming from the lower coverage in the subsampled data.

## Discussion

Copy number variations characterize different human disorders, and have special relevance in cancer^4,27^. In particular, tumors often present CNV heterogeneity, with several clones contained in the same tumor which may evolve differently during cancer progression and respond differently to treatment^1,28^. Measuring and quantifying the levels of CNV heterogeneity in cellular populations is therefore highly relevant^8^. Here, we present a novel computational method, epiAneufinder, which uses scATAC-seq data to faithfully recapitulate CNVs from single cells. This is achieved via segmenting the genome into equally sized bins and quantifying the number of reads mapping into every bin; to then test for significant differences in coverage depth along the genome. This is done by iteratively calculating the Anderson Darling (AD) distance between the read distributions on each side of every bin. The highest AD distances mark the positions of the most likely copy number variation breakpoints, based on which the genome is segmented. Every segment is then assigned to the state “normal”, “gain” or “loss” based on its mean read coverage. The cells are finally clustered based on the similarity of their gain and loss profiles, to identify CNV clones in the population.

Using an aneuploid cell line for which both scATAC-seq and scWGS data are available, we showed how the genome-wide CNV profiles recovered from scATAC-seq data by epiAneufinder compared to the ones obtained from scWGS, which can be considered a ground truth gold standard for the quantification of CNVs in single-cell populations. epiAneufinder was able to discover several clones in the population, harboring different CNV profiles containing both gains and losses.

Having established the performance of epiAneufinder compared to scWGS, we then moved to more physiologically relevant scenarios. Using two primary human samples of basal cell carcinoma that had been interrogated using scATAC-seq, we were able to identify multiple clones with distinct CNV profiles present in each tumor. This clonal variation could not be revealed by embedding and clustering of the same cells based on their genome-wide scATAC-seq peak profiles. These results highlight the relevance of studying single-cell heterogeneity based on individual copy number profiles, which is otherwise hidden in classical single-cell ATAC-seq analyses.

Finally, we performed a simulation to study the robustness of epiAneufinder copy number gain and loss calls to different sequencing depths. As expected, for lower sequencing depths copy number gain identification becomes more challenging, while more copy number losses are wrongly called. However, the overall copy number profiles are recovered across the genome even at 20% of the initial coverage.

In summary, epiAneufinder extracts single-cell copy number variations from scATAC-seq data alone, or alternatively from single-cell multiome data, without the need to supplement the data with other data modalities. The method leverages read depth distribution along the genome to infer a separate karyotype for every individual cell, allowing to explore the CNV heterogeneity present in a sample at the single-cell level. epiAneufinder unlocks a layer of genomic variation which is otherwise unexplored with traditional scATAC-seq data analysis, and its application offers the opportunity to explore relevant sources of heterogeneity which would otherwise remain hidden.

## Methods

### scATAC-seq and scWGS datasets

The scATAC-seq dataset for the SNU601 cell line^19^ was downloaded from the Short Read Archive (SRA) accession PRJNA674903, and the two pre-treatment basal cell carcinoma samples^18^ were obtained from GEO accession GSE129785 (accession number GSM3722057 for SU006 and GSM3722064 for SU008). The scWGS data for the SNU601 cell line^23^ was downloaded from SRA accession PRJNA498809. The scATAC data of the gastric cell line SNU601 were aligned to the human genome hg38 using the 10x Genomics Cell RangerAtac 2.0.0 software with default parameters. For the BCC samples, dataset was downloaded already aligned to the human genome version hg19.

### SNU601 CNV calling of scWGS data and comparison with scATAC

The scWGS of the SNU601 cell line^23^ was analyzed using aneufinder^24^ with default parameters and window size 100Kb. Aneufinder identified a cluster of cells with very high ploidy, cluster 2, consisting of 134 cells (SI Fig. 3), which we excluded from the scWGS data for the comparisons between scATAC and scWGS. Since the cell lines were measured in different laboratories, clonal variation among them can be expected^25^.

For the comparison between the two different data modalities, first we retained the same windows in both datasets. Afterwards a pseudo-bulk CNV profile was generated by counting the number of gain/loss/normal cells identified per window, and multiplying each gain by 3, each loss by 1 and each normal state by 2:

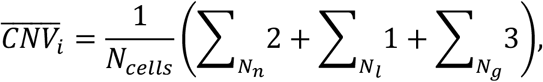

where *N*_*cells*_ is the total number of cells, *N*_*n*_ is the number of “normal” cells, *N*_*l*_ is the number of “loss” cells and *N*_*g*_ is the number of “gain” cells, at bin *i*. Pearson correlation and Mean Square Error (MSE), with 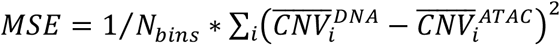, were calculated between the ATAC and the DNA profiles, both genome wide and per chromosome.

### WES data analysis

Whole exome sequencing data was downloaded^26^ from SRA accession PRJNA533341 (accession numbers SRS4645189 for SU006 tumor and SRS4645187 SU006 control; SRS4645177, SRS4645174 for SU008 tumor and SRS4645174 for SU008 control) (average 110-fold coverage) and aligned to the hg19 human genome using the Burrow-Wheeler aligner^29^ (version 0.7.17) with parameters mem -P -M -t32. Duplicates were removed (picard tools^30^ version 2.25.7) and base quality scores were recalibrated using GATK^31^ version 4.2.2., in accordance with Yost et al.^26^. To compare the results of epiAneufinder with the WES dataset, the number of WES reads within the respective epiAneufinder windows were calculated and tumor reads were normalized by the control reads. In addition, the windows were normalized by the percentage of exonic base pairs, and a gaussian filter was applied over the signal (the optimal sigma of the gaussian filter was chosen based on the correlation results between the pseudo-bulk and WES signals.). That signal was then correlated (Pearson correlation) to the pseudo-bulk scATAC-seq results, calculated as described in the section above for each sample (SI Fig. 6).

### Embedding of primary samples based on scATAC-seq profile

Peaks were called using MACS2^32^ on the aggregated dataset. The epiScanpy toolkit^33^ was used for subsequent analysis. First, we filtered observations to match the barcodes from the epiAneufinder output, and built a count matrix and computed gene activity. Then the peak matrix was binarized, variability scores (as defined by epiScanpy) were calculated per feature and the features were selected based on a variability score threshold of 0.53. Prior to further analysis the peak matrix was library size normalized and logarithmized. Principal component analysis was performed and the most informative PCs were selected using the elbow method (SU006: 5, SU008: 4). Using these top PCs we computed a neighborhood graph and an embedding using UMAP. The Leiden community detection algorithm was used for clustering and we identified the top differentially open peaks between clusters which were used for enrichment analysis with GREAT^34^ (SI Fig. 7). Moreover, several marker genes for tumor cells, fibroblasts and endothelial cells were quantified per cluster^18,35^ (SI FIg. 7). After evaluating marker gene activity and GO terms, clusters were annotated. We finally computed the composition of the cell type clusters with respect to the karyotypes, and vice versa.

### Downsampling of scATAC dataset and analysis

The SNU601 cell line dataset was downsampled with the 10x Genomics Cell Ranger Atac 2.0.0 software^18^ using the count command (subsample rate from 0.2 to 0.9). The results of the downsampling process can be viewed in Sup. Table 3. epiAneufinder was run for each percentage of the original dataset, with the same parameters as for the full set. In the filtering step of the algorithm, both cells and bins that did not pass the epiAneufinder quality controls were removed (Sup. Table 4). To compare the downsampled datasets with the full dataset, only common cells and bins were considered. For these, we quantified the number of bins that did not change their CNV status and the number of bins that changed their status from gain/loss/normal to a different one (Fig. 4).

To further explore the robustness of the algorithm we calculated “precision”, “recall” and “F1” scores using as a ground truth the CNV calls from the fully covered dataset (Sup. Table 5). For the gain state, we defined as true positive (*tp*) the bins with gain state in both the downsampled and the full set, as false positive (*fp*) the bins with gain in the downsampled but not in the full set and false negative (*fn*) the bins with gain in the full set but not in the downsampled one. Similarly we defined true positives, true negatives and false negatives for the loss and disomic states. Then the precision, recall and F1 scores were calculated as:

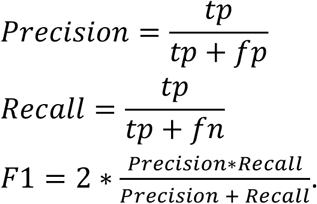

## Data availability

The following publically available datasets analyzed in this study were downloaded: The scATAC-seq dataset for the SNU601 cell line(Wu et al. 2021) was downloaded from the Short Read Archive (SRA) accession PRJNA674903, and the two pre-treatment basal cell carcinoma samples(Satpathy et al. 2019) were obtained from SRS accession GSE129785 (accession number GSM3722057 for SU006 and GSM3722064 for SU008). The scWGS data for the SNU601 cell line(Andor et al. 2020) was downloaded from SRA accession PRJNA498809.

## Code availability

epiAneufinder is available through Github (https://github.com/colomemaria/epiAneufinder).

## Acknowledgments

We would like to thank Antonio Scialdone for discussions about calling CNVs in scRNA-seq data, and Anna Danese for help in embedding of scATAC-seq data. This work was supported by the Impuls-und Vernetzungsfonds of the Helmholtz-Gemeinschaft (grant VH-NG-1219) for M.C.T. and A.R.. A.S. acknowledges support from HelmholtzAI. P.H. is supported by the Helmholtz Association under the joint research school “Munich School for Data Science - MUDS”.

## Contributions

MCT designed the study. MCT and AR conceived the algorithm. AR and AS implemented the algorithm. AR, AS and PH analyzed data. MS provided code. All authors contributed to manuscript writing.

## References

1. Bakhoum, S. F. & Landau, D. A. Chromosomal Instability as a Driver of Tumor Heterogeneity and Evolution. Cold Spring Harb. Perspect. Med. 7, (2017).

2. Taylor, A. M. et al. Genomic and Functional Approaches to Understanding Cancer Aneuploidy. Cancer Cell 33, 676–689.e3 (2018).

3. Sheltzer, J. M. & Amon, A. The aneuploidy paradox: costs and benefits of an incorrect karyotype. Trends Genet. 27, 446–453 (2011).

4. Ben-David, U. & Amon, A. Context is everything: aneuploidy in cancer. Nat. Rev. Genet. 21, 44–62 (2020).

5. Weaver, B. A. & Cleveland, D. W. The aneuploidy paradox in cell growth and tumorigenesis. Cancer cell vol. 14 431–433 (2008).

6. Shahbazi, M. N. et al. Developmental potential of aneuploid human embryos cultured beyond implantation. Nature Communications vol. 11 (2020).

7. Stankiewicz, P. & Lupski, J. R. Structural variation in the human genome and its role in disease. Annu. Rev. Med. 61, 437–455 (2010).

8. Bakker, B., van den Bos, H., Lansdorp, P. M. & Foijer, F. How to count chromosomes in a cell: An overview of current and novel technologies. Bioessays 37, 570–577 (2015).

9. Mallory, X. F., Edrisi, M., Navin, N. & Nakhleh, L. Methods for copy number aberration detection from single-cell DNA-sequencing data. Genome Biol. 21, 208 (2020).

10. Mallory, X. F., Edrisi, M., Navin, N. & Nakhleh, L. Assessing the performance of methods for copy number aberration detection from single-cell DNA sequencing data. PLOS Computational Biology vol. 16 e1008012 (2020).

11. Hou, Y. et al. Single-cell triple omics sequencing reveals genetic, epigenetic, and transcriptomic heterogeneity in hepatocellular carcinomas. Cell Res. 26, 304–319 (2016).

12. Serin Harmanci, A., Harmanci, A. O. & Zhou, X. CaSpER identifies and visualizes CNV events by integrative analysis of single-cell or bulk RNA-sequencing data. Nat. Commun. 11, 89 (2020).

13. Fan, J. et al. Linking transcriptional and genetic tumor heterogeneity through allele analysis of single-cell RNA-seq data. Genome Res. 28, 1217–1227 (2018).

14. Gao, R. et al. Delineating copy number and clonal substructure in human tumors from single-cell transcriptomes. Nat. Biotechnol. 39, 599–608 (2021).

15. Müller, S., Cho, A., Liu, S. J., Lim, D. A. & Diaz, A. CONICS integrates scRNA-seq with DNA sequencing to map gene expression to tumor sub-clones. Bioinformatics 34, 3217–3219 (2018).

16. Tickle, T., Tirosh, I., Georgescu, C., Brown, M. & Haas, B. inferCNV of the Trinity CTAT Project. Klarman Cell Observatory, Broad Institute of MIT and Harvard (2019).

17. Buenrostro, J. D., Giresi, P. G., Zaba, L. C., Chang, H. Y. & Greenleaf, W. J. Transposition of native chromatin for fast and sensitive epigenomic profiling of open chromatin, DNA-binding proteins and nucleosome position. Nat. Methods 10, 1213–1218 (2013).

18. Satpathy, A. T. et al. Massively parallel single-cell chromatin landscapes of human immune cell development and intratumoral T cell exhaustion. Nat. Biotechnol. 37, 925–936 (2019).

19. Wu, C.-Y. et al. Integrative single-cell analysis of allele-specific copy number alterations and chromatin accessibility in cancer. Nat. Biotechnol. (2021) doi:10.1038/s41587-021-00911-w.

20. Amemiya, H. M., Kundaje, A. & Boyle, A. P. The ENCODE Blacklist: Identification of Problematic Regions of the Genome. Sci. Rep. 9, 9354 (2019).

21. Anderson, T. W. & Darling, D. A. A Test of Goodness of Fit. J. Am. Stat. Assoc. 49, 765–769 (1954).

22. Pettitt, A. N. A two-sample Anderson-Darling rank statistic. Biometrika 63, 161–168 (1976).

23. Andor, N. et al. Joint single cell DNA-seq and RNA-seq of gastric cancer cell lines reveals rules of in vitro evolution. NAR Genom Bioinform 2, lqaa016 (2020).

24. Bakker, B. et al. Single-cell sequencing reveals karyotype heterogeneity in murine and human malignancies. Genome Biol. 17, 115 (2016).

25. Ben-David, U. et al. Genetic and transcriptional evolution alters cancer cell line drug response. Nature 560, 325–330 (2018).

26. Yost, K. E. et al. Clonal replacement of tumor-specific T cells following PD-1 blockade. Nat. Med. 25, 1251–1259 (2019).

27. Taylor, A. et al. MS12.02 Genomic and Functional Approaches to Understanding Cancer Aneuploidy. Journal of Thoracic Oncology vol. 14 S179 (2019).

28. Salgueiro, L. et al. Acquisition of chromosome instability is a mechanism to evade oncogene addiction. EMBO Mol. Med. 12, e10941 (2020).

29. Li, H. & Durbin, R. Fast and accurate short read alignment with Burrows–Wheeler transform. Bioinformatics 25, 1754–1760 (2009).

30. Picard. http://broadinstitute.github.io/picard/.

31. DePristo, M. A. et al. A framework for variation discovery and genotyping using next-generation DNA sequencing data. Nat. Genet. 43, 491–498 (2011).

32. Zhang, Y. et al. Model-based analysis of ChIP-Seq (MACS). Genome Biol. 9, R137 (2008).

33. Danese, A. et al. epiScanpy: integrated single-cell epigenomic analysis. Nature Communications 12, 5228 (2021).

34. McLean, C. Y. et al. GREAT improves functional interpretation of cis-regulatory regions. Nat. Biotechnol. 28, 495–501 (2010).

35. Franzén, O., Gan, L.-M. & Björkegren, J. L. M. PanglaoDB: a web server for exploration of mouse and human single-cell RNA sequencing data. Database 2019, (2019).

