## Supplementary figures and tables for "epiAneufinder: identifying copy number variations from single-cell ATAC-seq data"

### Supplementary Figures and Tables: epiAneufinder: identifying copy number variations from single-cell ATAC-seq data

**Figure S1: Number of reads distribution per somy**

a)

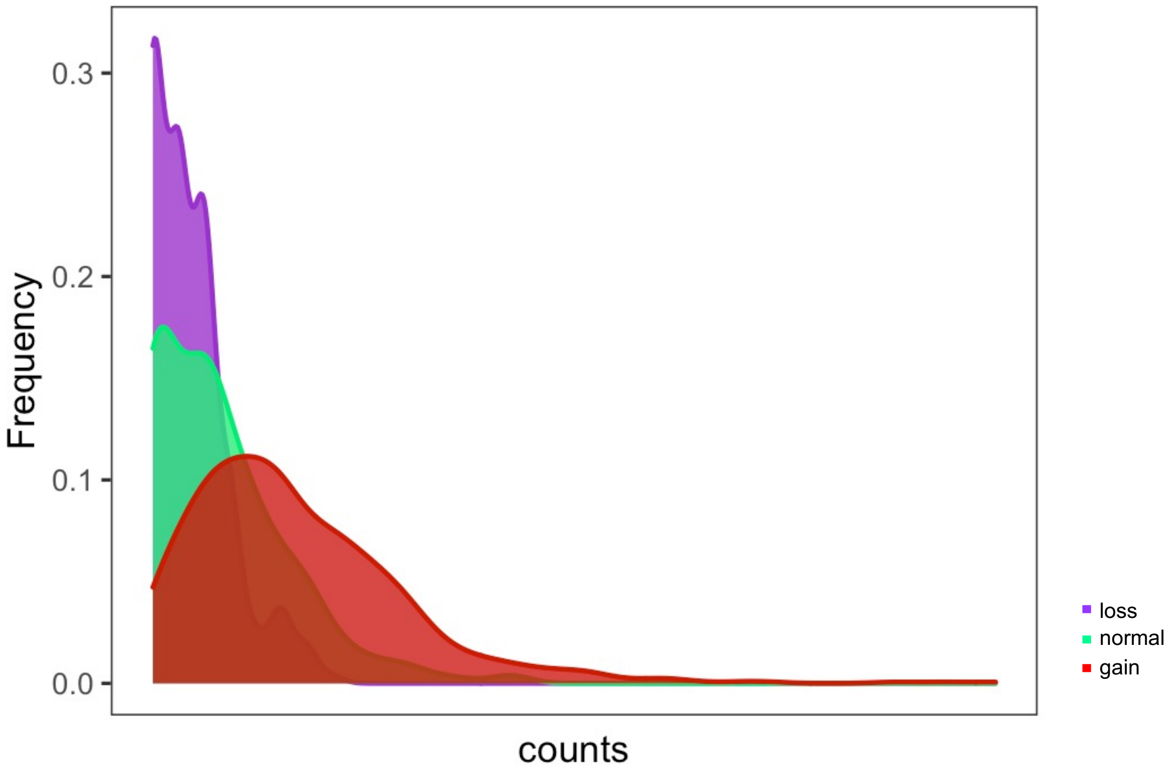


b)

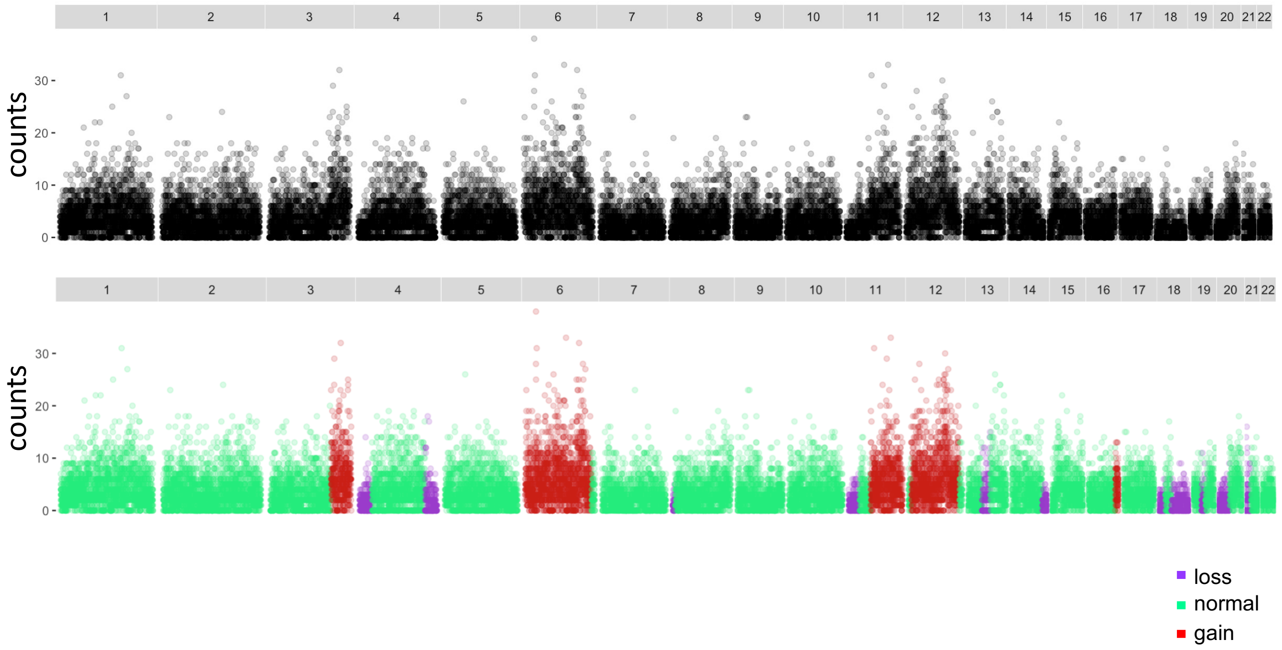


**Figure S1:** a) Histogram of the number of reads mapping into every 100 000 bp bin, for the regions identified as lost (purple), gained (red) and normal (green), showing how the distributions are different per somy. b) genome-wide representation (chromosome 1 to 22) of the number of counts per 100 000 bp bin for the same cell, with copy numbers indicated by purple (loss), red (gain) and green (normal). The data belongs to one single-cell from the SNU601 cell line (Wu et al. 2021), and the copy numbers were identified using epiAneufinder.

#### Figure S2: SNU segmentation with clustering results for scATAC-seq

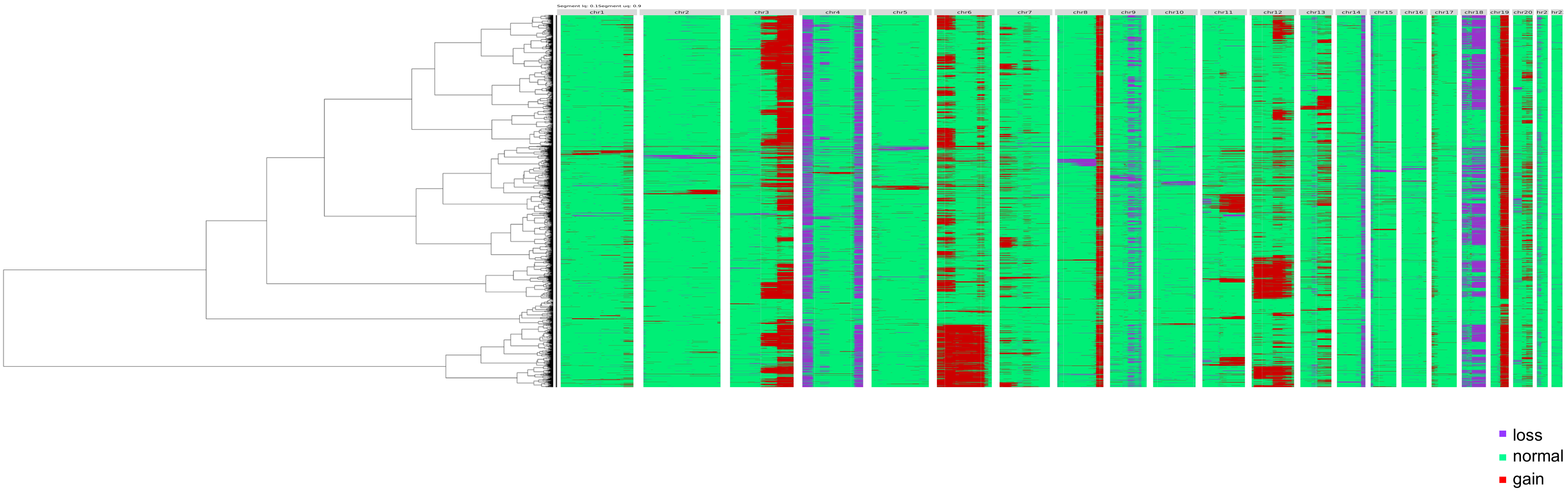


**Figure S2:** Karyogram per single-cell (every cell is a row, every column is a chromosome) for the SNU601 scATAC-seq dataset (Wu et al. 2021), with gains indicated in red, losses in pruple and normal state in green. The cells are ordered based on the clustering results (clustering based on their copy number profiles using Euclidean distance and Ward Clustering).

#### Figure S3: SNU601 segmentation for WGS

a)

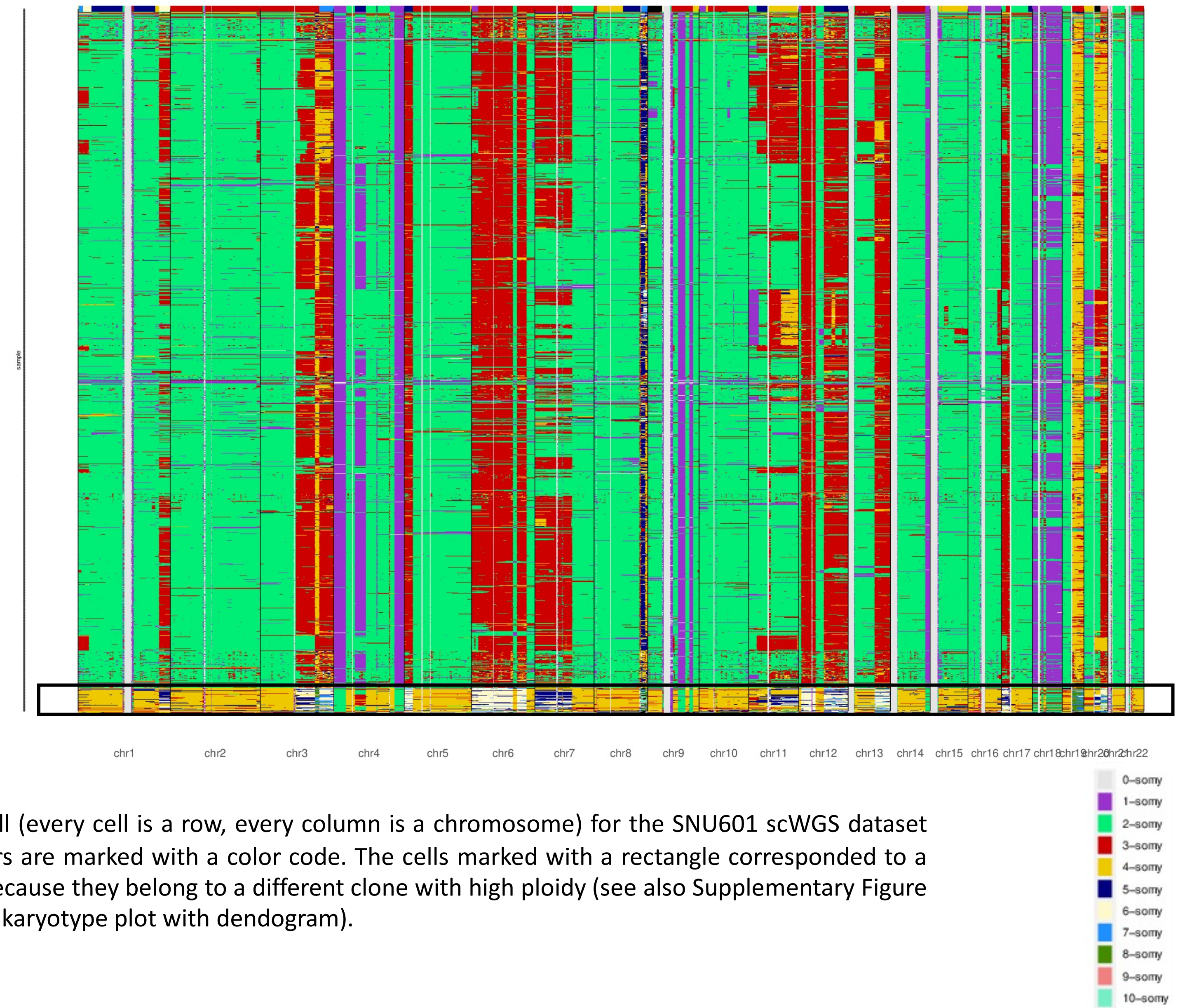


**Figure S3:** a) Karyogram per single-cell (every cell is a row, every column is a chromosome) for the SNU601 scWGS dataset (Andor et al. 2020). The copy numbers are marked with a color code. The cells marked with a rectangle corresponded to a separate cluster and were removed because they belong to a different clone with high ploidy (see also Supplementary Figure 3b on a separate file for the extended karyotype plot with dendogram).

#### Figure S4: SU006 patient sample segmentation with clustering results for scATAC-seq

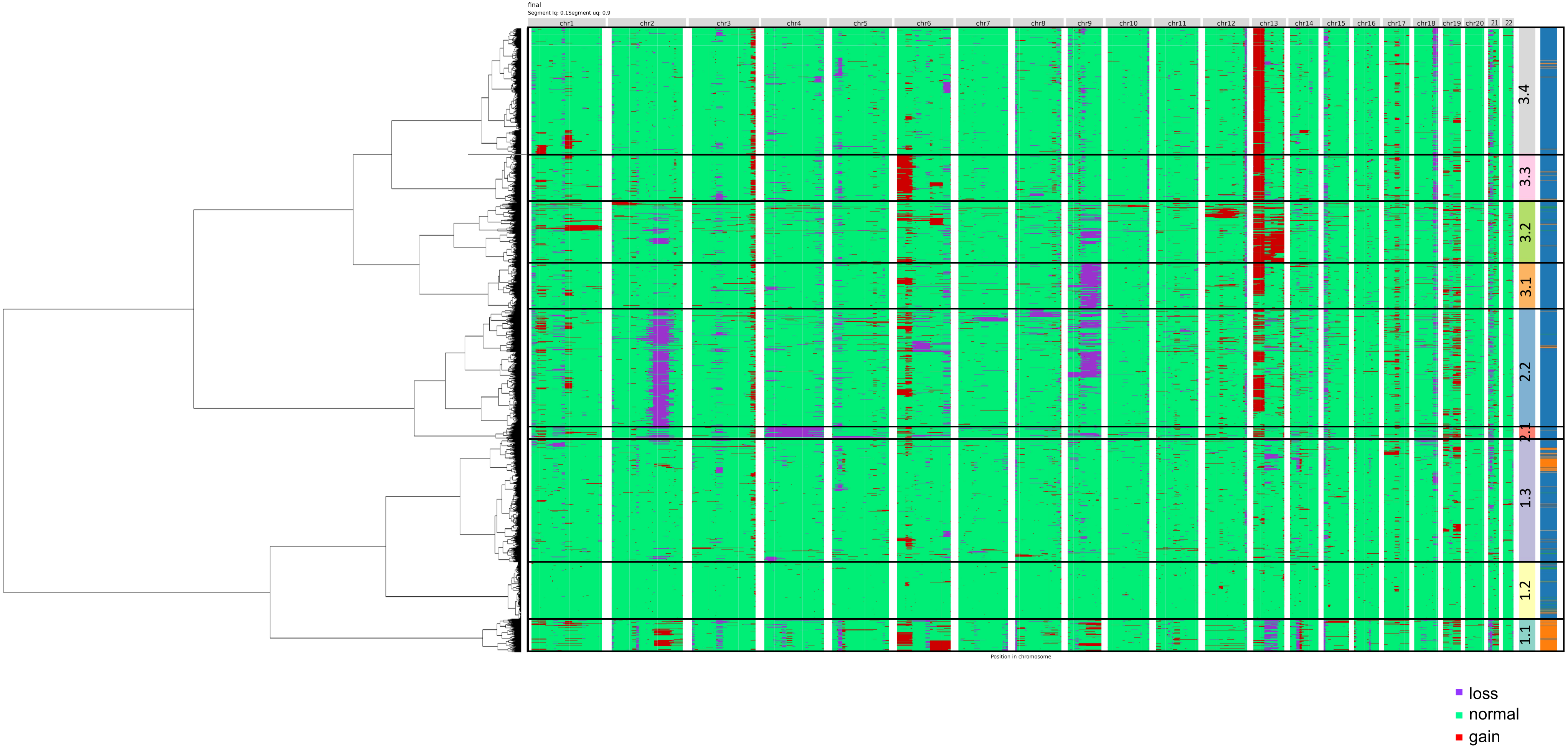


**Figure S4:** Karyogram per single-cell (every cell is a row, every column is a chromosome) for the SU006 patient sample (Satpathy et al. 2019), with gains indicated in red, losses in pruple and normal state in green. The cells are ordered based on the clustering results (clustering based on their copy number profiles using Euclidean distance and Ward Clustering). Clones are indicated for branch cutting at depth 9. The colour bar represents cell types from SI figure 7.

#### Figure S5: SU008 patient sample segmentation with clustering results for scATAC-seq

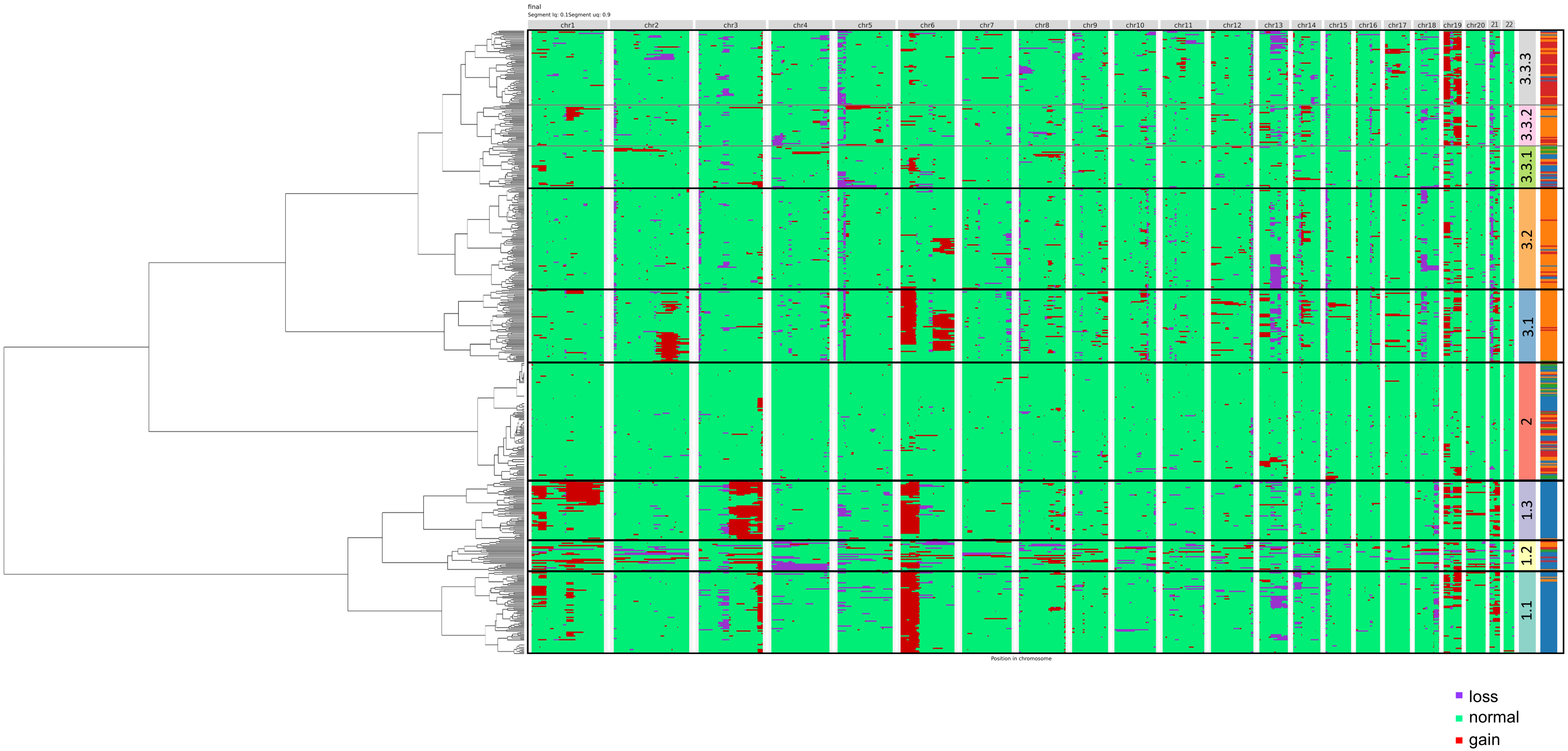


**Figure S5:** Karyogram per single-cell (every cell is a row, every column is a chromosome) for the SU008 patient sample (Satpathy et al. 2019), with gains indicated in red, losses in pruple and normal state in green. The cells are ordered based on the clustering results (clustering based on their copy number profiles using Euclidean distance and Ward Clustering). Clones are indicated for branch cutting at depth 7. The colour bar represents cell types from SI figure 7.

#### Figure S6: Comparison of WES to scATAC-seq for primary patient data

a)

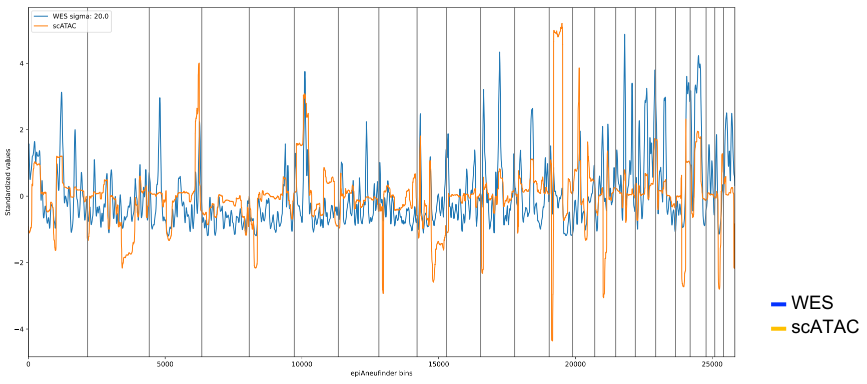


b)

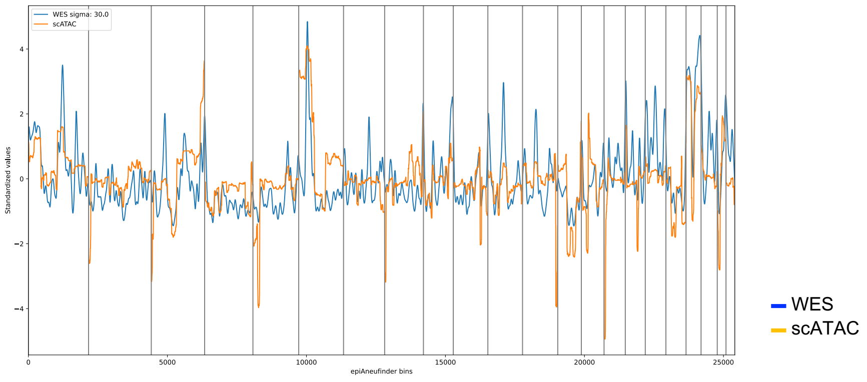


**Figure S6:** Comparison of WES (Yost et al. 2019 ) copy number profiles to aggregated (pseudo bulk) scATAC-seq profiles (Satpathy et al. 2019) for patients SU006 (a) and SU008 (b).

**Figure S7: SU006 and SU008 patient sample embedding based on scATAC-seq**

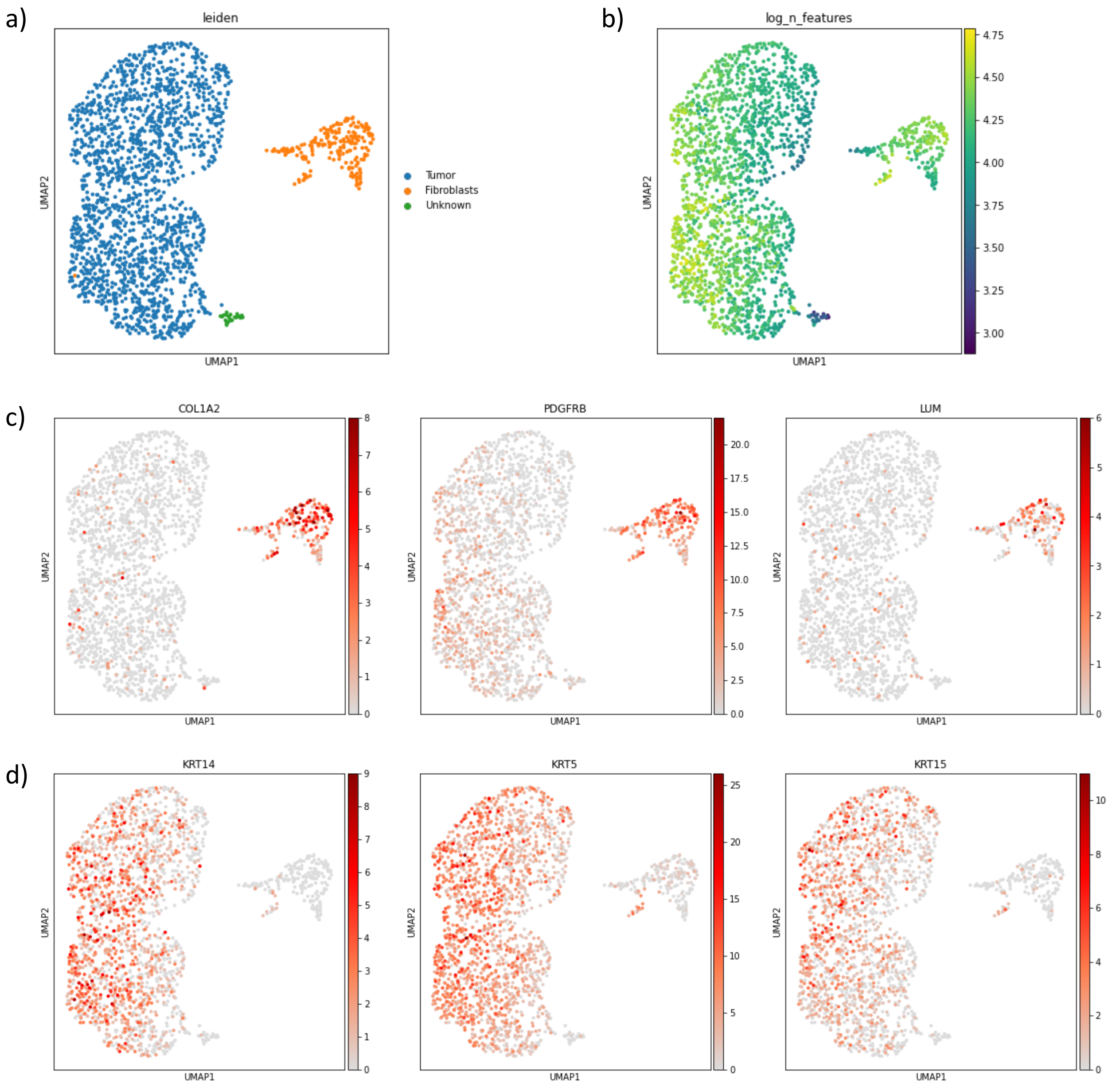


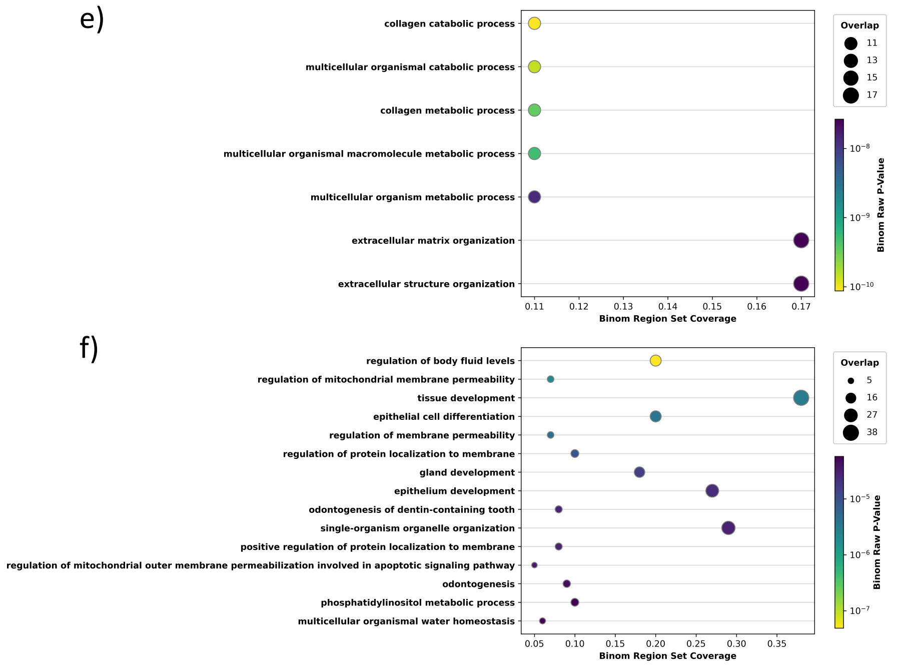


**Figure S7: Embedding based on scATAC-seq data for patients SU006 and SU008** (Satpathy et al. 2019). a) SU006: Cell types are indicated. b) SU006: number of features, where it can be observed that the “unknown” cluster has a low number of features. c) SU006: Gene activity levels for fibroblast genes (COL1A2, PDGFRB and LUM). d) SU006: Gene activity levels for basal cell carcinoma tumor marker genes (KRT14, KRT5, KRT15). e-f) SU006: GO terms for fibroblasts (e) and tumor cells (f).

**Figure S7: SU006 and SU008 patient sample embedding based on scATAC-seq**

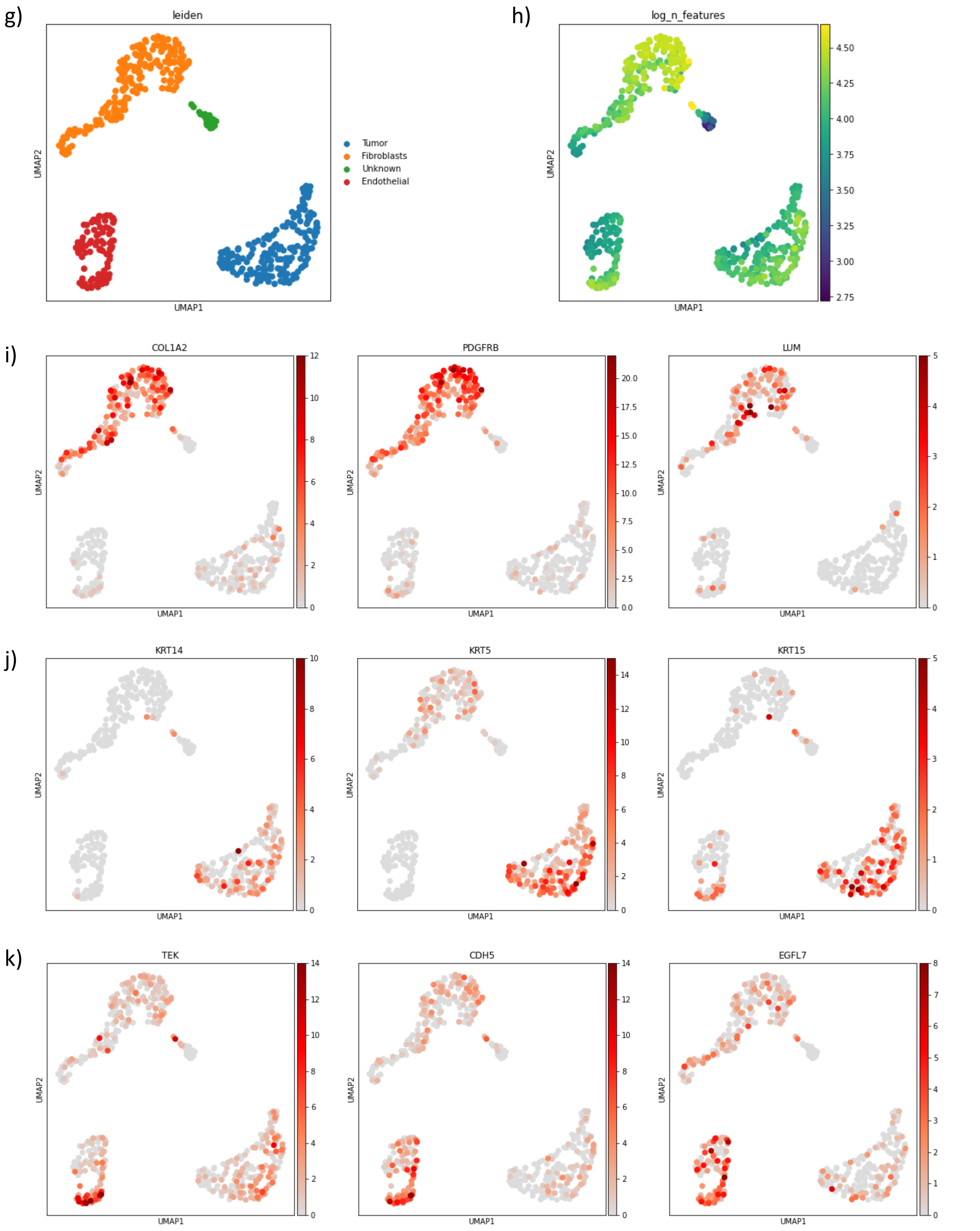


**Figure S7: Embedding based on scATAC-seq data for patients SU006 and SU008** (Satpathy et al. 2019). g) SU008: Cell types are indicated. h) SU008: number of features, where it can be observed that the "unknown" cluster is characterized by a low number of features. i) SU008: Gene activity levels for fibroblast genes (COL1A2, PDGFRB and LUM). j) SU008: Gene activity levels for basal cell carcinoma tumor marker genes (KRT14, KRT5, KRT15). k) SU008: Gene activity levels for endothelial cell marker genes (TEK, CDH5, EGFL7).

#### Figure S7: SU006 and SU008 patient sample embedding based on scATAC-seq

l)

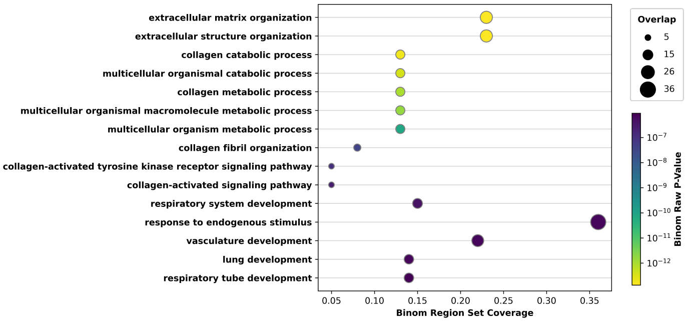


m)

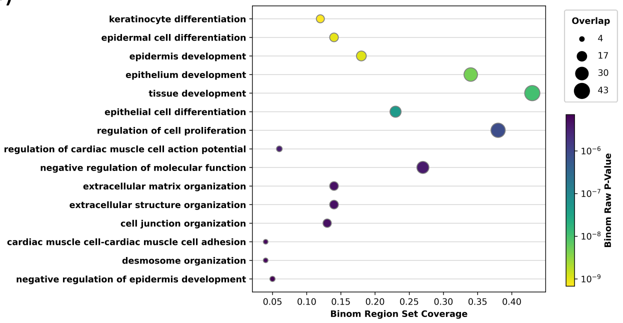


n)

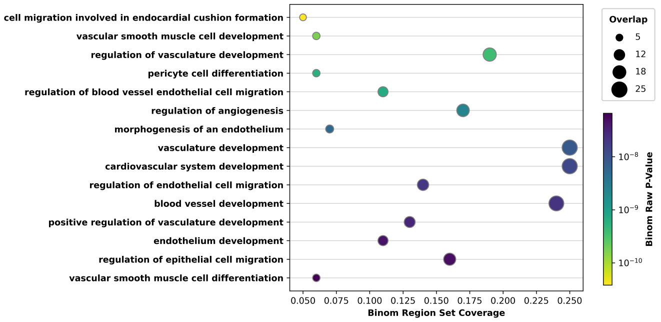


**Figure S7: Embedding based on scATAC-seq data for patients SU006 and SU008** (Satpathy et al. 2019). l-n) GO terms for fibroblasts (l), tumor cells (m) and endothelial cells (n).

#### Figure S8: Leiden clusters versus karyotype clones in patient samples

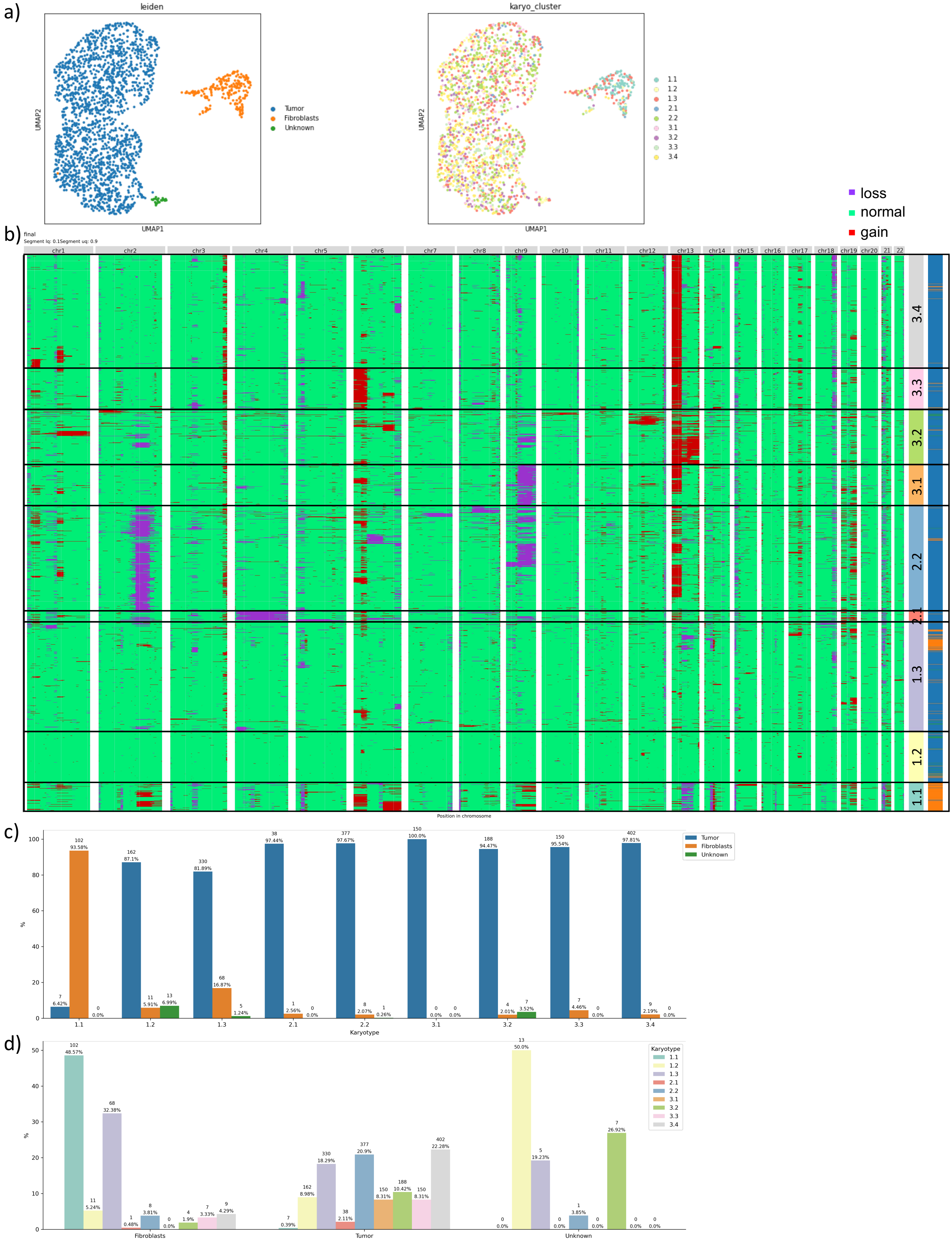


**Figure S8: Patient SU006** (Satpathy et al. 2019) a) Embedding based on scATAC-seq data with cell types (left) and karyotype clones (right). b) Karyotype profile with indicated karyotype clones and corresponding cell types. c) Correspondence between the cells in every karyotype clone and Leiden cluster. d) Correspondence between the cells in every Leiden cluster and karyotype clone.

#### Figure S8: Leiden clusters versus karyotype clones in patient samples

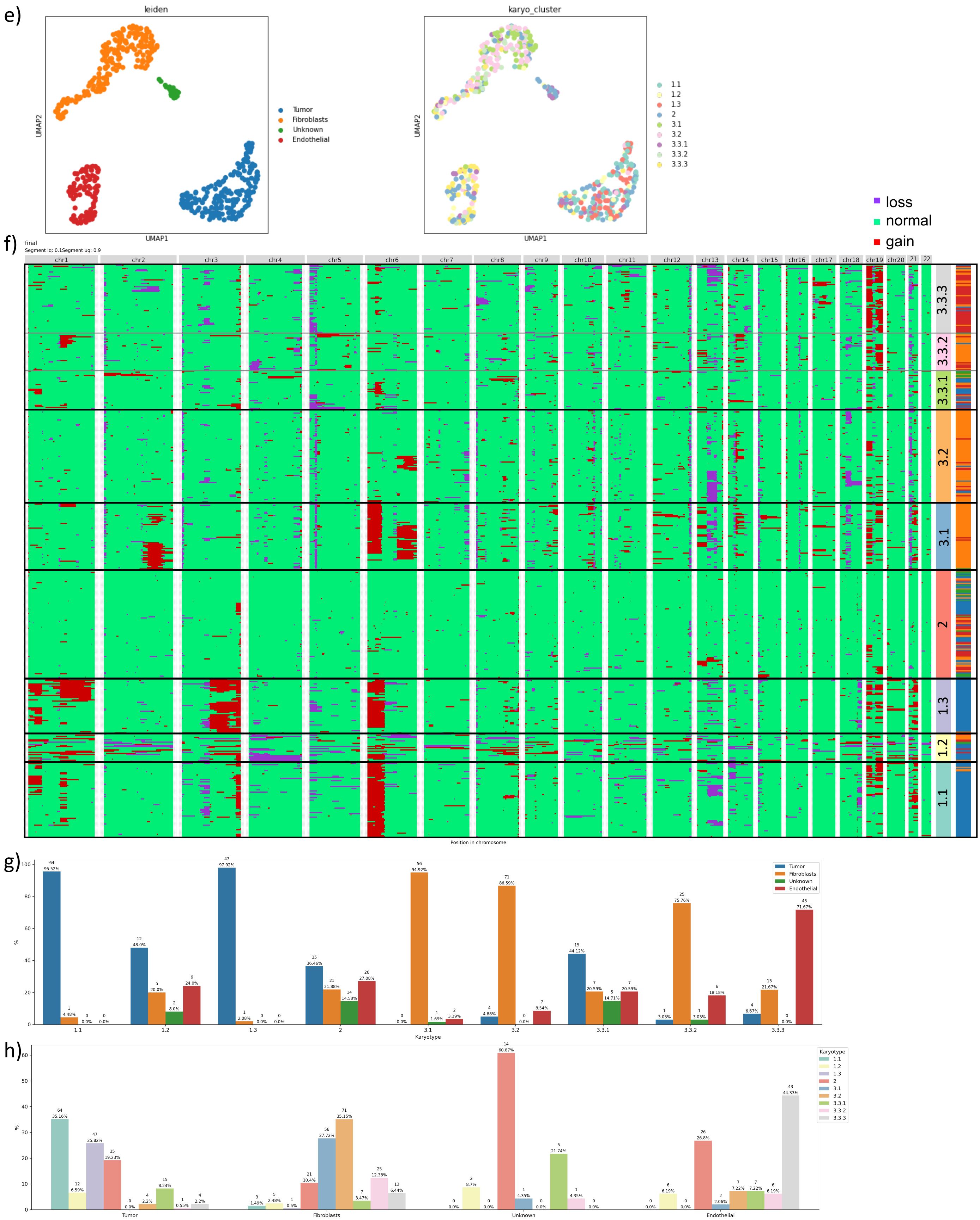


**Figure S8: Patient SU008** (Satpathy et al. 2019) e) Embedding based on scATAC-seq data with cell types (left) and karyotype clones (right). f) Karyotype profile with indicated karyotype clones and corresponding cell types. g) Correspondence between the cells in every karyotype clone and Leiden cluster. h) Correspondence between the cells in every Leiden cluster and karyotype clone.

**Table S1: Runtime**

| Sample | Number of cells | RAM Usage | CPU wall time |
|---|---|---|---|
| SNU601 100% | 3157 | 160.16 GB | 16:51:56 |
| SNU601 90% | 3123 | 141.47 GB | 17:59:21 |
| SNU601 80% | 3089 | 137.87 GB | 17:00:09 |
| SNU601 70% | 3043 | 123.66 GB | 15:53:38 |
| SNU601 60% | 2977 | 147.69 GB | 14:35:07 |
| SNU601 50% | 2883 | 144.96 GB | 13:57:15 |
| SNU601 40% | 2748 | 105.67 GB | 12:18:37 |
| SNU601 30% | 2489 | 86.81 GB | 10:06:38 |
| SNU601 20% | 1908 | 64.48 GB | 07:33:15 |
| SU008 Tumor | 504 | 41.55 GB | 04:22:03 |
| SU006 Tumor | 2040 | 83.97 GB | 11:35:10 |

**Sup. Table 1 :** CPU wall time and memory usage of different datasets. For all datasets a single node in the cluster was used, with 6 cores and 192Gb RAM. For the wall time calculations we used all the steps of the epiAneufinder pipeline.

**Table S2:**

| | Mean Square Error |
|---|---|
| genome | 0,088302155 |
| chr1 | 0,044066379 |
| chr2 | 0,002747019 |
| chr3 | 0,05630817 |
| chr4 | 0,061856663 |
| chr5 | 0,102142792 |
| chr6 | 0,272204255 |
| chr7 | 0,140853308 |
| chr8 | 0,006562585 |
| chr9 | 0,128786441 |
| chr10 | 0,00227922 |
| chr11 | 0,057532808 |
| chr12 | 0,290062742 |

| | Mean Square Error |
|---|---|
| chr13 | 0,212218036 |
| chr14 | 0,01587572 |
| chr15 | 0,008588208 |
| chr16 | 0,008908706 |
| chr17 | 0,120135691 |
| chr18 | 0,087198595 |
| chr19 | 0,087600032 |
| chr20 | 0,172552848 |
| chr21 | 0,031805875 |
| chr22 | 0,004677832 |

**Sup. Table 2:** Mean square error for the comparison between the copy number calls from scWGS and scATAC-seq.

**Table S3:**

| SNU601 Percent coverage | cells | fragments per cell | Mean read pairs per cell | Sequenced read pairs |
|---|---|---|---|---|
| 1 | 3575 | 75013 | 147358.63 | 526807107 |
| 0.9 | 3547 | 70117 | 132921.31 | 474130324 |
| 0.8 | 3555 | 64738 | 118548.79 | 421440958 |
| 0.7 | 3541 | 58944 | 104138.51 | 368754462 |
| 0.6 | 3534 | 52679 | 89439.96 | 316080807 |
| 0.5 | 3527 | 45843 | 74679.96 | 263396204 |
| 0.4 | 3528 | 38420 | 59725.61 | 210711941 |
| 0.3 | 3502 | 30298 | 45124.65 | 158026510 |
| 0.2 | 3462 | 21323 | 30430.35 | 105349877 |

**Sup. Table 3:** Results from downsampling the SNU601 cell line dataset.

**Table S4:**

| SNU601 Percent coverage | cells after filtering | bins after filtering |
|---|---|---|
| 1 | 3154 | 26560 |
| 0.9 | 3086 | 26543 |
| 0.8 | 3086 | 26543 |
| 0.7 | 3040 | 26526 |
| 0.6 | 2974 | 26512 |
| 0.5 | 2880 | 26498 |
| 0.4 | 2475 | 26481 |
| 0.3 | 2486 | 26441 |
| 0.2 | 1908 | 26310 |

**Sup. Table 4:** Number of cells and bins that were retained after the filtering step of epiAneufinder for the different downsampling percentages.

**Table S5:**

| | Precision | | | Recall | | | F1 | | |
|---|---|---|---|---|---|---|---|---|---|
| Percent coverage | Gain | Loss | Disomic | Gain | Loss | Disomic | Gain | Loss | Disomic |
| 0.2 | 0.797823 | 0.637100 | 0.936620 | 0.606774 | 0.713194 | 0.954952 | 0.689305 | 0.673003 | 0.945697 |
| 0.3 | 0.805474 | 0.694934 | 0.946569 | 0.666197 | 0.764596 | 0.958414 | 0.729245 | 0.728102 | 0.952454 |
| 0.4 | 0.817965 | 0.737720 | 0.953865 | 0.716054 | 0.786472 | 0.962699 | 0.763624 | 0.761316 | 0.958261 |
| 0.5 | 0.831866 | 0.768682 | 0.958447 | 0.746264 | 0.803911 | 0.966286 | 0.786743 | 0.785902 | 0.962351 |
| 0.6 | 0.837794 | 0.793247 | 0.963371 | 0.776435 | 0.827105 | 0.968247 | 0.805948 | 0.809822 | 0.965803 |
| 0.7 | 0.851394 | 0.818158 | 0.967805 | 0.808224 | 0.840223 | 0.971347 | 0.829248 | 0.829044 | 0.969572 |
| 0.8 | 0.865776 | 0.844223 | 0.972655 | 0.841337 | 0.858143 | 0.974527 | 0.853382 | 0.851126 | 0.973590 |
| 0.9 | 0.890533 | 0.883269 | 0.978566 | 0.875845 | 0.888536 | 0.979917 | 0.883127 | 0.885894 | 0.976542 |

**Sup. Table 5:** Precision, recall and F1 score of the gain/loss/disomic bins in the different downsampled fractions
