## Supplementary figures and images for "epiAneufinder: identifying copy number variations from single-cell ATAC-seq data"

### Supplementary figure 3b

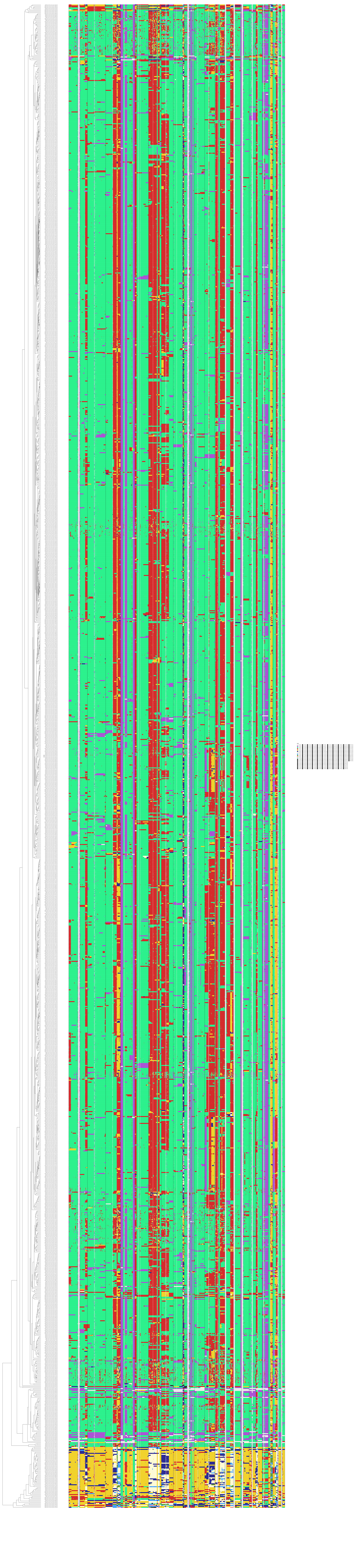
